# Hericerin derivatives from *Hericium erinaceus* exert BDNF-like neurotrophic activity in central hippocampal neurons and enhance memory

**DOI:** 10.1101/2020.08.28.271676

**Authors:** Ramón Martínez-Mármol, YeJin Chai, Zahra Khan, Seon Beom Kim, Seong Min Hong, Rachel S. Gormal, Dae Hee Lee, Jae Kang Lee, Mi Kyeong Lee, Sun Yeou Kim, Frédéric A. Meunier

## Abstract

The traditional medicinal mushroom *Hericium erinaceus* has long been known for enhancing the peripheral nerve regeneration through targeting nerve growth factor (NGF) neurotrophic activity. It was also reported to protect against ageing-dependent cognitive decline in wildtype and in Alzheimer’s disease mouse models suggesting a yet to be defined action on neurons of the central nervous system. Here, we purified and identified biologically active compounds from *H. erinaceus*, based on their ability to promote neurite outgrowth in hippocampal neurons. *N*-de phenylethyl isohericerin (NDPIH), an isoindoline compound from this mushroom together with its hydrophobic derivative hericene A, were highly potent in inducing extensive axon outgrowth and neurite branching in the absence of serum demonstrating high neurotropic activity. NDPIH also induced enlarged growth cones suggestive of a brain-derived neurotrophic factor (BDNF)-like activity. Pharmacological inhibition of tropomyosin receptor kinase B (TrkB) by ANA12 prevented NDPIH-induced neurotrophic activity providing evidence that NDPIH acts *via* TrkB receptors to mediate its neurotrophic effect in central neurons. Finally, *in vivo* treatment with *H. erinaceus* crude extract and hericene A significantly increased BDNF and downstream pathway and enhanced learning and memory in the novel object recognition memory test. Our results suggest that hericene A can promote BDNF-like activity in neurons *in vitro* and *in vivo* thereby enhancing recognition memory.

## Introduction

Neurotrophins are a family of molecules that promote neuronal survival, neurite outgrowth, and dendritic branching through binding specificity for particular tropomyosin receptor kinase (Trk) high-affinity receptors and p75 low-affinity receptors (Kaplan & Miller 2000). Mammalian neurotrophins comprises nerve growth factor (NGF), brain-derived neurotrophic factor (Ip *et al*. 1993) (BDNF), neurotrophin 3 (NT3) and neurotrophin 4 (NT4; also known as NT5) (Chao 2003; Ip *et al*. 1993). NGF, for example, can induce growth cone turning and sprouting of axonal filopodia through binding to TrkA receptor (Gallo & Letourneau 1998). BDNF induces the formation of filopodia and lamellipodia in growth cones from chick sensory neurons and from Xenopus spinal neurons *via* binding to TrkB receptor (Gallo & Letourneau 1998; Gibney & Zheng 2003). Both NGF and BDNF neurotrophins also bind to p75 receptor and prevent cell death. BDNF in particular, is highly expressed in the adult central nervous system and is critically important for the survival of neurons located in areas of the brain involved in memory acquisition as such as the hippocampus, and the cortex (Allen *et al*. 2011). Not surprisingly, BDNF plays a critical role in synaptic plasticity, and the BDNF receptor TrkB has been shown to be essential for long term potentiation, an Hebbian mechanism strengthening synaptic output in response to a high frequency train of stimulation, as well as learning (Allen *et al*. 2011). Dysfunction of the BDNF pathway has been linked with several disorders of the brain, including Alzheimer’s disease, schizophrenia (Xiu *et al*. 2009), Huntington’s disease (Zuccato & Cattaneo 2009), and Rett syndrome (Zuccato & Cattaneo 2009). Reduced BDNF mRNA and protein levels were found in the hippocampus and cortex of Alzheimer’s disease patients (Allen *et al*. 2011). The neurotrophin hypothesis poses that dysfunctions of neurotrophin pathways drive the pathogenic process for these disease (Appel 1981). For all these reasons, neurotrophins such as BDNF have attracted much attention for their neurotrophic and neuroprotective properties in central nervous system and have been used for therapeutic treatment of a range of neurodegenerative disorders including Alzheimer’s disease. BDNF treatment led to protection of forebrain cholinergic neurons (Knusel *et al*. 1992) and was associated with a reduction in amyloid beta (Aβ) in rat brains (Arancibia *et al*. 2008) and in nonhuman primates (Nagahara *et al*. 2009). Similarly, BDNF infusion provided significant neuronal protection in a Parkinsonian nonhuman primate model following treatment with the dopaminergic neurotoxin 1-methyl-4-phenyl-1,2,3,6-tetrahydropyridine (MPTP) (Deng *et al*. 2016).

However, translation of BDNF exogenous treatments into the clinic have failed initial scrutiny for various reasons including short half-life, poor blood–brain barrier (BBB) permeability, and off-target effects (Chan *et al*. 2017). Alternative interventions to increase BDNF levels have emerged including sustained drug-delivery system, engineered cell delivery and biomaterials providing more effective access to the brain (Houlton *et al*. 2019). Raising endogenous BDNF levels through exercise and botanical remedy is also an area of great interest to attempt to counteract the devastating effect of neurodegeneration.

For these reasons, ongoing efforts are made to identify compounds derived from natural sources. Traditional Chinese medicine provides with an enticing source of botanical remedy as several common drugs have stemmed from the Chinese pharmacopeia and been used for millennia throughout Asia and India for their therapeutic or poisonous pharmacologic profiles. One of the most promising nootropic fungi known for its neurotrophic profile comes from a *Hericium erinaceus*, also known as Lion’s Mane mushroom, which, paradoxically, was used to treat unrelated ailments such as stomach aches and as prophylactic treatment of cancers (Kim *et al*. 2013). Several studies have reported a strong neurotrophic effect along with the identification of numerous bioactive components, including polysaccharides, erinacines, hericerins, alkaloids, steroids, and many others (Friedman 2015; Brandalise *et al*. 2017). Interestingly, hericenones and erinacines have been shown to effectively cross the BBB (Hu *et al*. 2019) and proven to have neuroprotective effect both *in vitro* and *in vivo* in animal models of peripheral nerve injury (Wong *et al*. 2012), stroke (Hazekawa *et al*. 2010) and Alzheimer’s disease (Friedman 2015; Mori *et al*. 2011; Kawagishi & Zhuang 2008). This suggested a dual action on the peripheral and central nervous system.

Compounds derived from *H. erinaceus* have been shown to promote NGF synthesis and secretion (Zhang *et al*. 2015; Mori *et al*. 2009; Lai *et al*. 2013; Raman *et al*. 2015; Kawagishi *et al*. 1996) and to act via the TrkA pathway involving MEK/ERK and PI3K/Akt activation (Haure-Mirande *et al*. 2017; Phan *et al*. 2014). These neuroprotective effects were revealed in neurons derived from the peripheral nervous system (Ustun & Ayhan 2019). More recently, *H. erinaceus* was suggested to also affects the neurotrophin BDNF pathway in 1321N1 human astrocytoma cell line (Rupcic *et al*. 2018), and in mice brains (Chiu *et al*. 2018). It was also shown to increase circulating pro-BDNF levels in humans (Vigna *et al*. 2019), suggesting a general effect on BDNF synthesis and on TrkB pathway via an unknown mechanism.

We purified and identified several compounds for their ability to promote an efficacious neurotrophic effect on cultured hippocampal neurons fully capable of replacing foetal calf serum, critically involved in hippocampal neuron survival in culture. Neurite outgrowth and branching are increase by several fold and super-resolution structured illumination microscopy revealed much enlarged growth cone morphology in treated neurons. This neurotropic effect was partially blocked by ANA-12, a well-characterised inhibitor of TrkB activation (Cazorla *et al*. 2011). We further found that dietary supplementation with *H. erinaceus* crude extract significantly enhanced recognition memory. Importantly, supplementation with 20-50 times less concentration of purified hericene A was equally potent enhancing recognition memory. Tested rodents exhibited elevated BDNF and TrkB downstream effectors levels. Overall, our results demonstrate that hericene A is a memory enhancer and act via the BDNF/TrkB pathway.

## Materials and Methods

### Preparation of samples

The fruiting bodies of *H. erinaceus* were provided from the C&G Agricultural Association (Sejong, Korea). Column chromatography was performed on silica gel (Kieselgel 60, 70–230 mesh, Merck, Darmstadt, Germany), and semi-preparative HPLC was performed using a Waters HPLC system (Waters, Milford, MA, USA) equipped with two Waters 515 pumps with a 2996 photodiode-array detector.

The dried fruiting body of *H. erinaceum* was extracted with EtOH at room temperature and filtered with filter paper, which yielded the ethanol extract (A1) and filter cake. The ethanol extract was suspended in H_2_O and partitioned with CH_2_Cl_2,_ EtOAc, *n*-BuOH, successively, to yield the CH_2_Cl_2_-soluble (A2), EtOAc-soluble (A3), *n*-BuOH soluble (A4) and water fraction (A5). The filter cake was further extracted with water to yield water extract (A6).

The CH_2_Cl_2_-soluble fraction (A2) was chromatographed on silica gel eluting with a mixture of *n*-hexane-EtOAc by step gradient to give 12 subfractions (A2A-A2L). C1 and C3 were obtained from A2F and A2K, respectively, by semi-preparative HPLC eluting with MeCN-water (45:55). C2 and C4 were obtained from A2D by recrystallization followed by semi-preparative HPLC eluting with MeCN. The structures of isolated compounds were identified as *N*-de phenylethyl isohericerin (C1), hericene A (C2), corallocin A (C3) and 4-[3′,7′′-dimethyl-2′,6′-octadienyl]-2-formyl-3-hydroxy-5-methyoxybenzylalcohol (C4) by the analysis of the spectroscopic data and comparison with literature values (Li *et al*. 2015; Wittstein *et al*. 2016).

### Reagents

Extracts from Lion’s mane mushroom and Ginkgo leaves were weight and dissolved in dimethyl sulfoxide (DMSO) to obtain stock solutions concentrated at 0.1, 1 and 10 mg/ml. Working solutions of the extracts were prepared by diluting 1:1,000 the stocks in neurobasal medium at the following concentrations 0 µg/ml, 0.1 µg/ml, and 10 µg/ml. Hippocampal neurons at 2DIV or 3DIV were incubated with the extracts during 24 h. After the treatments (3DIV or 4DIV), neurons were prepared for immunocytochemistry. Recombinant human BDNF and ANA-12 (Sigma-Aldrich) was incubated during 24h at 1 nM and 50 μM, respectively.

### Primary hippocampal cultures

Hippocampal neurons were obtained from Sprague Dawley (SD) rats at embryonic day (E)18 and prepared as described previously (Joensuu *et al*. 2017; Joensuu *et al*. 2016). Hippocampal neurons used in Figs. 1-3, were seeded at 20,000 onto 35mm poly-L-lysine-coated glass-bottom dishes (Celvis) or coverslips in Neurobasal growth medium supplemented with 2% B27, 2 mM Glutamax, 50 U/ml penicillin, 50 μg/ml streptomycin, and 5% fetal bovine serum (FBS) (Wang *et al*. 2020; Wang *et al*. 2016; Wang *et al*. 2015). At DIV1 (1 day in vitro), neurons were switched to serum-free medium until use at DIV4.

**Figure 1.**
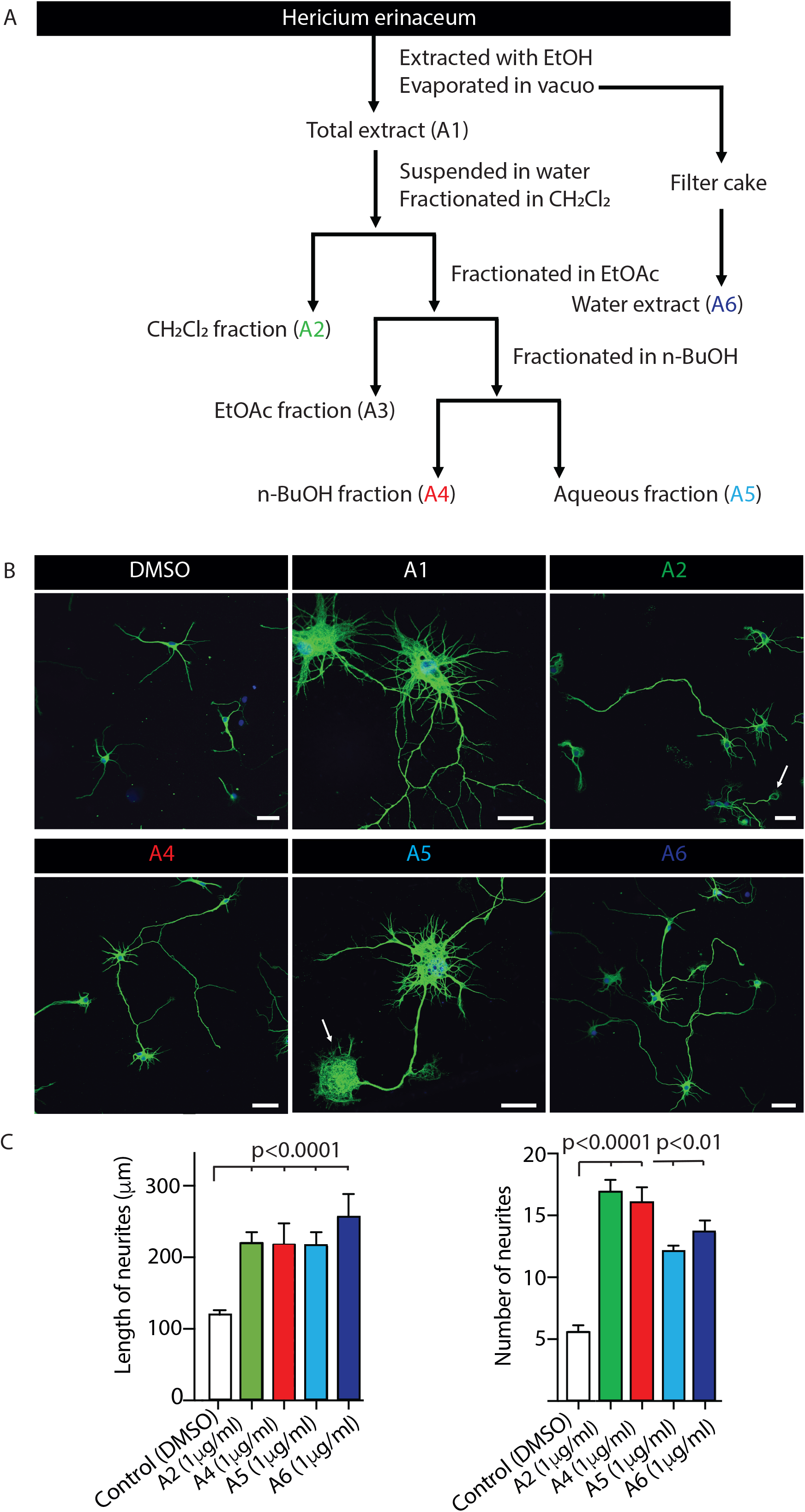
Lion Mane mushroom (LMM) extracts exert a potent neurotrophic effect in hippocampal neurons. (A) Scheme of the purification procedure to generate tested extracts (A1-A6). Hippocampal neurons were cultured in the presence of FBS (5%) for 24h, the starved until treated with indicated extracts at DIV3. (B) Representative images of hippocampal neurons (DIV3) treated for 24h with control vehicle (DMSO), or indicated extracts (A1-A6) purified from Lion Mane mushroom. Neurons were then fixed and processed for immunofluorescence against β-tubulin (green) and imaged using confocal microscopy (nuclear DAPI in blue). (C) Longest neurite (axon) length and numbers quantification for each indicated treatments. Data shows mean ± SEM, n=30-60 neurons in each condition, from 3 independent neuronal preparations. One-way ANOVA, Turkey’s multiple comparison test was performed. P-value is indicated when significative differences were found. Scale bar 100 µm.

Hippocampal neurons used in Figs. 4-5, were seeded at lower density (10,000) onto a poly-L-lysine-coated dish in the complete absence of serum to avoid any interference and paracrine effect. The neurons were cultured in Neurobasal medium (Gibco) supplemented with 2% B27 (Gibco), 2 mM Glutamax (Gibco) and 50 U/mL penicillin/ streptomycin (Invitrogen) for 2 days *in vitro* (2DIV) after being treated with specific reagents. All animal procedures were approved by the University of Queensland Animal Ethics Committee.

### Immunocytochemistry

Primary hippocampal neurons were fixed with, 4% paraformaldehyde/4% sucrose in PBS for 10 min at room temperature, and permeabilized with, 0.1% Triton X-100 in PBS for 10 min. To detect the actin cytoskeleton, neurons were stained with a solution of 1 µg/mL Alexa-Fluor™ 647-phalloidin (Thermo Fisher) in PBS for 60 minutes and rinsed with PBS, 3 times, 5 minutes each. Neurons were blocked for 1 h in PBS, 5% goat serum, 0.025% Triton X-100, followed by primary antibody incubation overnight at 4 °C and secondary antibody incubation for 1 h at room temperature. The primary antibody used was mouse Anti-IIIβ-tubulin (MMS-435P, Covance, 1:1,000), and the secondary antibody was Alexa Fluor™ 488-labeled donkey-anti-mouse or Alexa Fluor™ 568-labeled donkey-anti-mouse (all from LifeTechnologies, Thermo Fisher, 1:500). Coverslips were mounted in ProLong medium (Invitrogen, Thermo Fisher). Fluorescence images were acquired with a 20x (0.8 NA / 550µm WD / 0.22 μm/pixel), or 63x (1.4 NA / 190µm WD / 0.19 μm/pixel) objective on a Zeiss LSM510 META confocal microscope. Structured Illumination Microscopy (SIM) was performed on a Zeiss PS1 ELYRA equipped with a 63× (1.4 NA / Plan-Apochromat / 190µm working distance / 0.064 μm/pixel/ Oil – For SIM) objective and a PCO scientific sCMOS camera. Images were acquired with an exposure time of 200 ms, a SIM grating size of 28 μm, using three rotations and 5 phases. Structured illumination images were then processed using Zen software.

### Image Analysis and Quantifications

Each experiment contains a mixed culture of neurons isolated from more than three embryos. All experiments were repeated three independent times (independent dissections). Between 30 to 60 randomly selected neurons were analyzed for each condition. For the analysis of axon length, the longest neurite of each neuron was selected. Neurite Tracer tool from FIJI-ImageJ (Schindelin *et al*. 2012) software was used to quantify the length of the axons, the number of neurites and growth cone area.

### Behavior tests of animals and sample administration

Male ICR mice (23 – 25 g), 6 weeks old, were purchased from Orient Bio Co. (Seoul, Korea), and fed on a laboratory diet (AIN-76A purified diet) and given water *ad libitum*. Mice were housed for 1 week at 23 ± 1°C with 55 ± 5% humidity under a 12 h light/dark cycle. The Gachon University Lab Animal Care Committee approved the care and use of the mice for the study (GIACUC-R2019001) in accordance with the Korean Guide for Care and Use of Laboratory Animals. The mice were randomly divided into four groups (5 mice per group) as followed: Vehicle group, piracetam group (positive control, PC) and A1 group. The normal (10 ml/kg of distilled water, per os, *p*.*o*.), PC (400 mg/kg, p.o.) or A1 (100 and 250 mg/kg, *p*.*o*.) or A2-C2 (5 mg/kg, *p*.*o*.) groups received oral administration of the substances, and each sample were dissolved in distilled water. Behavioral tests (*i*.*e*., Y-maze test, novel object recognition test) were performed on days 21-23 and mice were sacrificed on day 24 for biochemical analysis.

### Y-maze test

The Y-shaped maze consisted of three identical arms (60 × 15 × 12 cm^3^) made of dark grey plastic with equal angles between each arm. The three arms of Y shape were marked as A, B and C. Firstly, mice were placed into center of a Y-shaped runway and allowed to move throughout the maze for 8 min, and the move sequence (*e*.*g*., ABCCBA) and total number of arm entries was recorded manually (Bae et al., 2020). These parameters calculated included number of arm entries, same arm returns, alternate arm returns. The percentage of spontaneous alternation performance (%) was determined using equation (1). Equation (1) = [Actual alternations (total alternations)] / [Possible alternation (total number of arm entries – 2)] x 100.

### Novel object recognition task (NORT) test

The NORT test was performed by using an open box (60 × 60 × 60 cm^3^) (Mesripour et al., 2016). Each mouse was place in the open box with two different kind of identical objects for three minutes at the 1^st^ day for training. In the 2^nd^ day, each mouse was placed again in the box in which one of the identical objects had been replaced with a novel object. These parameters calculated included the time (in seconds) spent to explore the familiar object, time (in seconds) spent to explore the novel object, and total time (in seconds) spent to explore both objects. The percentage of recognition (%) was determined using equation (2). Equation (2) = [Time (in seconds) spent to explore the novel object] / [total time (in seconds) spent to explore both objects] x 100.

### Western blot analysis

Western blot assay was performed as previously described (Subedi et al., 2019). The total proteins (30 μg) which obtained from each brain tissue were separated by 10% acrylamide SDS-PAGE gel electrophoresis, and then transferred to PVDF membranes. The transferred PVDF membranes were incubated with primary antibodies including α-Tubulin, glyceraldehyde-3-phophate dehydrogenase (GAPDH), brain-derived neurotrophic factor (BDNF), nerve growth factor (NGF), glial cell-derived neurotrophic factor (GDNF), growth associated protein (GAP-43), cAMP-response element binding protein (CREB), phosphorylated (p)-CREB, extracellular-signal-regulated kinase (ERK) and p-ERK (1:1000, Santa Cruz Biotechnology, Inc., Santa Cruz, CA) at 4°C overnight. On the 2^nd^ day, the membranes were incubated with secondary antibodies, and protein bands were visualized using ECL Western Blotting Detection Reagents (Amersham Pharmacia Biotech, Little Chalfont, UK). Densitometric analysis was performed using Bio-Rad Quantity One software version 4.3.0 (Bio-Rad Laboratories, Inc.).

### Statistical Analysis

All the values shown in the graphs represent the mean ± SEM. The number of neurons and the statistical test used in each experiment are specified in the corresponding figure legends. Statistical tests were performed and figures were made using GraphPad Prism 7 software. The D’Agostino and Pearson test was used to test for normality of acquired data. Two-tailed, unpaired Student’s t-test was used to compare two conditions, and one-way ANOVA with Tukey post hoc test was used when the experiment compared more than two groups. Significance is considered when *p-value < 0.05, **p < 0.01, or ***p < 0.001.

## Results

To test the potential neurotrophic effect of *Hericium erinaceum* extracts, we performed various fractionations as indicated in Fig. 1A starting with an ethanol extract (A1), which was subsequently dried, resuspended in water and fractionated with dichloromethane (CH_2_Cl_2_) (A2) or further with ethyl acetate (EtOAc) (A3). The latter was further fractionated in *n*-butanol (*n*-BuOH) (A4) and in water (A5). Polysaccharide fraction (A6) was prepared from the filter cake after A1 was filtered. We dissolved all the fractions in DMSO and tested their effects on cultured hippocampal neurons. To carry out these experiments, hippocampal neurons from E18 rat were plated at relatively low density (20,000 cells per coverslip) and cultured in the presence of FBS (5%) for 24h. We then removed the serum and incubated them until DIV3, when we treated them with the extracts for 24 h prior to fixation and immunostaining processing with β-tubulin antibody (to visualise microtubules) and 4′,6-diamidino-2-phenylindole (DAPI, nuclear marker). We first checked that the vehicle DMSO used for the extracts did not affect the axonal growth and neurite formation, which was indeed the case (Fig. 1B, C). We also tested several unrelated extracts (from other mushrooms and Ginseng) at the same concentration and found no neurotrophic effect (data not shown). We next tested the effect of the Lion’s mane mushroom A2 extracts (1 μg/ml). Incubation with A2 lead to a clear neurotrophic effect (Fig. 1B) with axonal length increased by two-fold (Fig. 1C). The number of neurites detected was also greatly increased by more than 3-fold (Fig. 1C).

The high neurotrophic activity elicited by the A2 compounds suggested enhanced growth cone activity leading to increase of total axonal length and neurite formation. In fact, we also observed large growth cones in A2-treated hippocampal neurons. We next tested additional Lion’s mane mushroom compounds (A4-A6) extracted with an alternative extraction method (Fig. 1A). Interestingly, incubation with all extracts triggered significant neurotrophic effect detectable at concentrations of 1 μg/ml (Fig. 1B), with axonal length increasing by several-fold (Fig. 1C). The number of neurites detected was also greatly increased (Fig. 1C). The chloroform extract was further fractionated by chromatography to further separate several molecules determined by NMR (supp Fig. 1).

We identified *N*-de phenylethyl isohericerin (C1, NDPIH), hericene A (C2), corallocin A (C3), and 4-[3′,7′-dimethyl-2′,6′-octadienyl]-2-formyl-3-hydroxy-5-methyoxybenzylalcohol (C4) (Fig. 2A). All of these purified extracts had significant neurotrophic activity as indicated by the 3-fold increase in neurite length (Fig. 2B and 2C). As most extracts were active, we investigated their effects on the axonal growth cone molecular organization and architecture. We incubated the extracts A2 to A5, as well as NDPIH (C1) on hippocampal neurons for 24h and co-stained them using beta-tubulin and phalloidin-rhodamine (Fig. 3). Vehicle treated neurons exhibit classical growth cone architecture with actin localized in peripheral filopodia and tubulin located in the central (C) domain (Fig. 3A). A2, A4, A5 as well as NDPIH-treated neurons exhibited highly enlarged growth cones displaying arrays of large filopodia in the peripheral (P) domain, some of them larger than the cell body itself (Fig. 3A). Analysis of the area occupied by growth cones relative to DMSO-treated neurons reveal a 2-to 5-fold average increase in size (Fig 3B). We used structured illumination microscopy to assess whether the β-tubulin organization was affected by NDPIH treatment. We could find that most microtubule were present in the C-domain of growth cones and NDPIH treatment promoted formation of microtubule-enriched filopodial structures that extend along the P-domain of growth cones (Fig. 3C). The overall cytoskeletal architecture of growth cones was therefore preserved but the size was markedly increased suggestive of high neurotropic activity.

**Figure 2.**
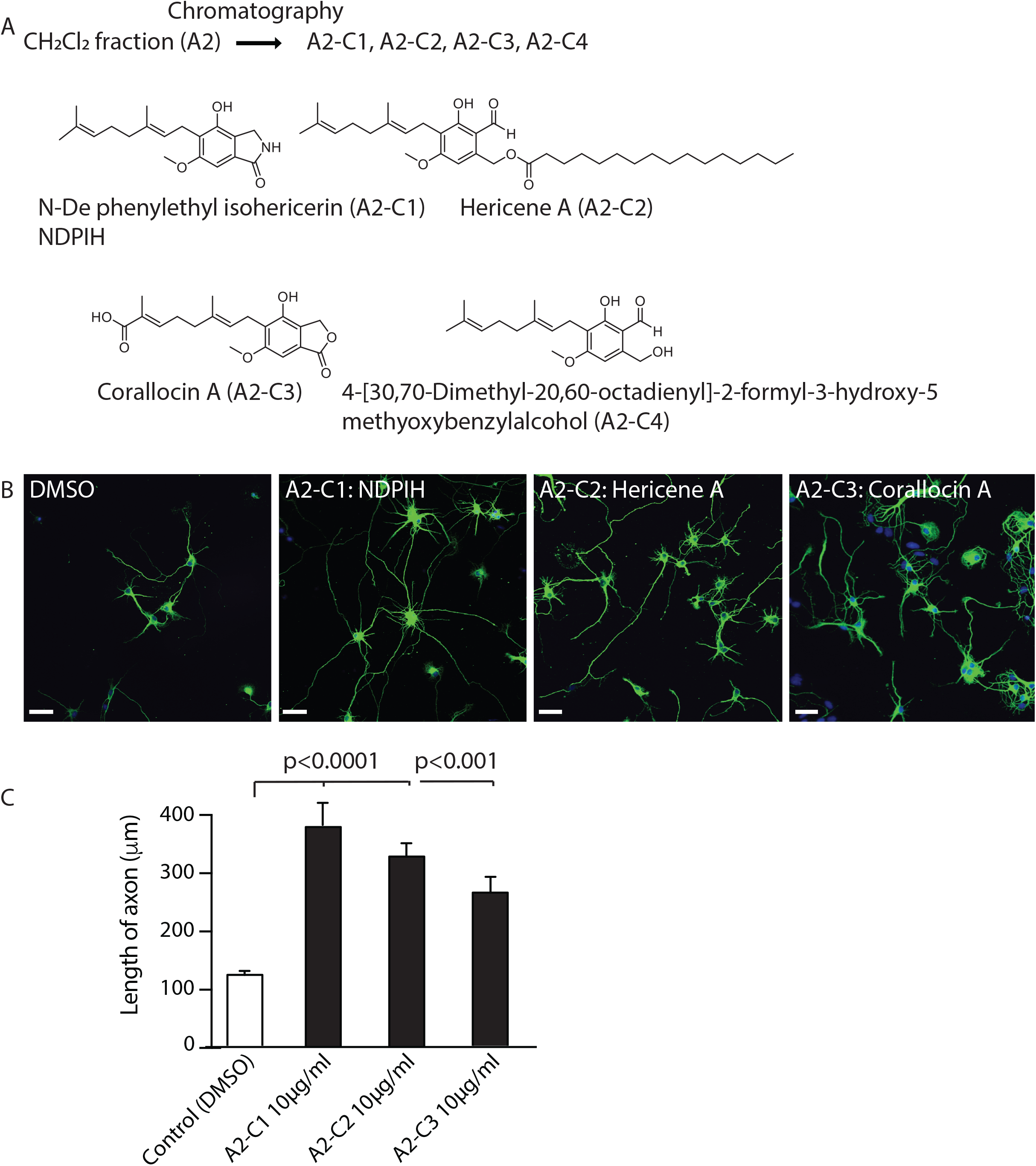
A2-derived compounds exert a potent neurotrophic effect in hippocampal neurons. (A) Chemical structures of identified A2-derived compounds. Hippocampal neurons were cultured in the presence of FBS (5%) for 24h, the starved until treated with indicated extracts at DIV3. (B) Representative images of hippocampal neurons (DIV3) treated for 24h with control vehicle (DMSO), or indicated extracts (NDPIH, Hericene A and Corallocin A). Neurons were then fixed and processed for immunofluorescence against β-tubulin (green) and imaged using confocal microscopy (nuclear DAPI in blue). (C) Longest neurite (axon) length quantification for each indicated treatments. Data shows mean ± SEM, n=30-60 neurons in each condition, from 3 independent neuronal preparations. Two-tailed, unpaired Student’s t-test was performed. P-value is indicated when significative differences were found. Scale bar 50 µm.

**Figure 3.**
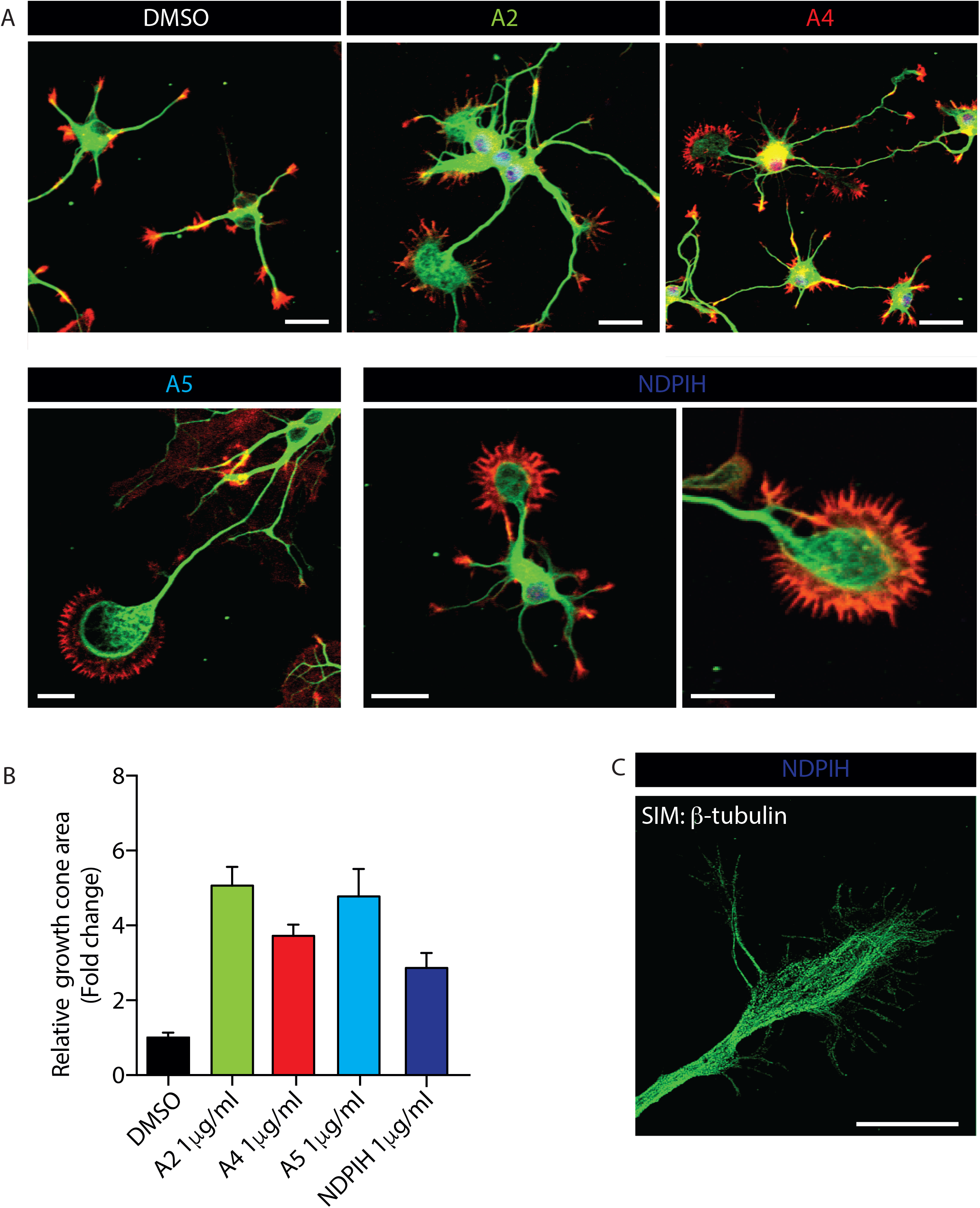
Extracts purified from Lion Mane mushroom (LMM) exert a neurotrophic effect on growth cone. Hippocampal neurons were cultured in the presence of FBS (5%) for 24h, the starved until treated with indicated extracts at DIV3. (A) Representative images of hippocampal neurons (DIV3) treated for 24h with control vehicle (DMSO), or indicated extracts (A2, A4, A5 and NDPIH) purified from Lion Mane mushroom. Neurons were then fixed and processed for immunofluorescence against β-tubulin (green) and actin filaments (red) and imaged using confocal microscopy (nuclear DAPI in blue). (C) Growth cone area quantification for each indicated treatments. Data shows mean ± SEM, n=15 neurons in each condition, from 4 independent neuronal preparations. Scale bar 50 µm. (C), Representative image of a NDPIH-treated neuron acquired using structure illumination microscopy. Note the classical distribution microtubules in this enlarged growth cone.

Overall, the *H. erinaceum* extracts were surprisingly active in hippocampal neurons with hericene A and NDPIH displaying high neurotrophic activity. However, it was not clear whether the compounds were capable to promoting this neurotropic activity directly or via an autocrine mechanism similar to that suggested for the NGF activity in peripheral neurons (Phan *et al*. 2014; Zhang *et al*. 2015). To investigate the ability of NDPIH to directly promote neurite outgrowth, we reduced further the number of neurons seeded to 10,000 per coverslip and omitted any serum during their culture. We therefore tested the ability of the extracts to directly promote neurite outgrowth. Neurons were therefore seeded at low density to prevent paracrine effects from neighbour cells and were exposed to NDPIH (Fig. 4A) at DIV2 for 24h. Neurons were then fixed and we processed for immunocytochemistry using anti-β-tubulin and DAPI (Fig. 4B). Importantly, NDPIH significantly increased neurite length in a concentration-dependent manner, strongly suggesting a direct BDNF-like neurotropic action in these central neurons (Fig. 4C).

**Figure 4.**
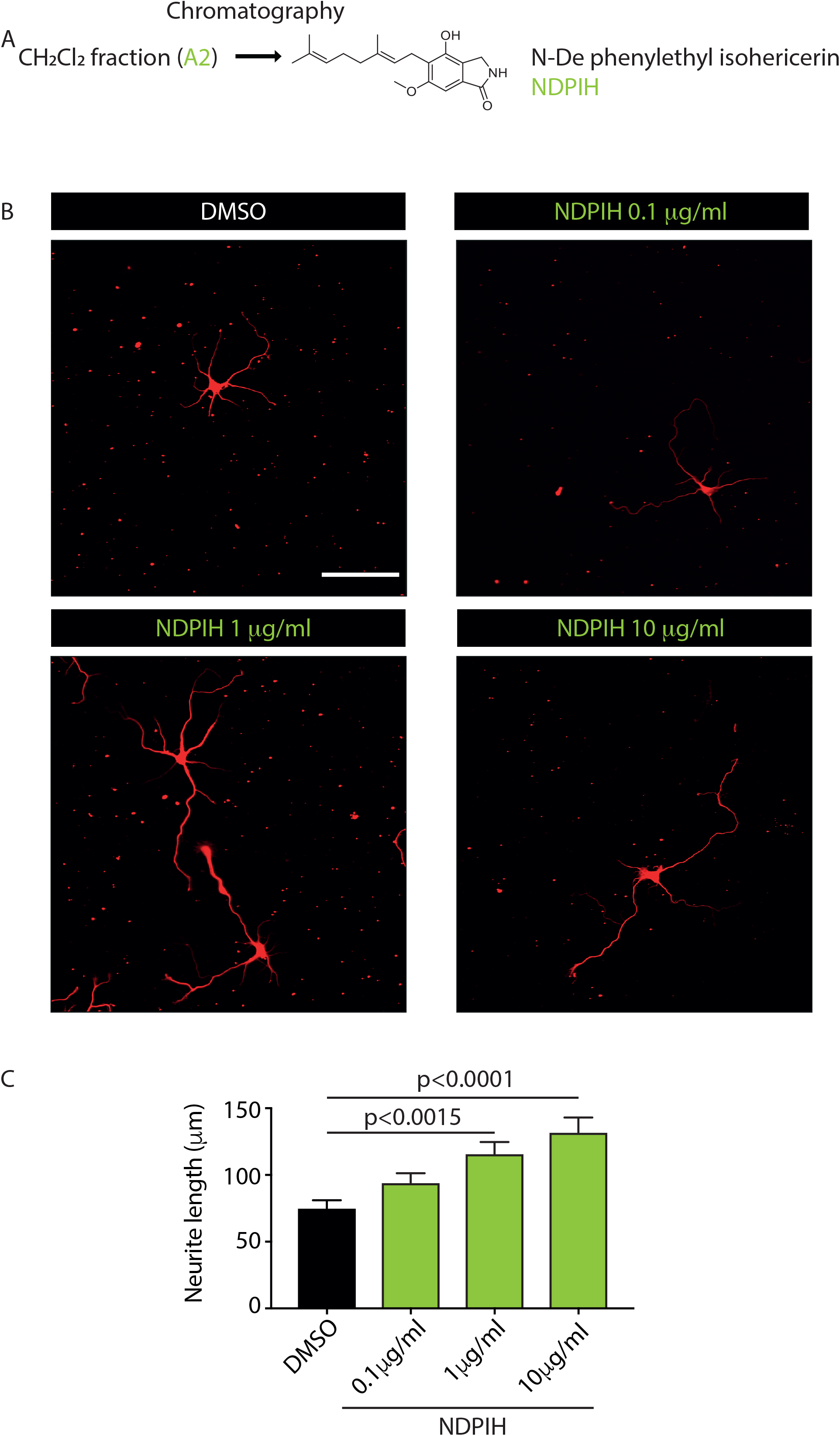
NDPIH purified from Lion Mane mushroom exerts a dose-dependent neurotrophic effect in hippocampal neurons. (A) Chemical structures of identified A2-derived compound NDPIH. Hippocampal neurons seeded at low density (10,000 per coverslip) to prevent paracrine effects until DIV2. Cells were then exposed to indicated concentration of purified NDPIH for 24h, fixed, processed for immunochemistry against β-tubulin (green) and imaged using confocal microscopy (nuclear DAPI in blue). (B) Representative images of hippocampal neurons, treated with control vehicle (DMSO) or indicated concentrations (0.1, 1 or 10 μg/ml) of NDPIH purified from Lion Mane mushroom extract A2. (C) Axon length quantification for each treatment. Axon was considered as the longest neurite. Data shows mean ± SEM. n=30-60 neurons in each condition, from 3 independent neuronal preparations. One-way ANOVA, Turkey’s multiple comparison test was performed. P-value is indicated when significative differences were found. Scale bar 100 µm.

To test the hypothesis that NDPIH act via a BDNF pathway, we incubated neurons with the BDNF inhibitor ANA-12, a specific TrkB antagonist capable of selectively preventing TrkB auto-phosphorylation by BDNF (Cazorla *et al*. 2011). As anticipated, co-incubation the mushroom extract with ANA-12 (0.5 μM) significantly reduced the neurotropic activity-induced by NDPIH (Fig. 5A-B). These results confirmed that, at least, some of the observed NDPIH neurite-outgrowth activities were directly through the TrkB pathway. Importantly, only half of the effect by NDPIH was inhibited by ANA-12 (Fig. 5C), suggesting that additional pathways may also be involved.

**Figure 5.**
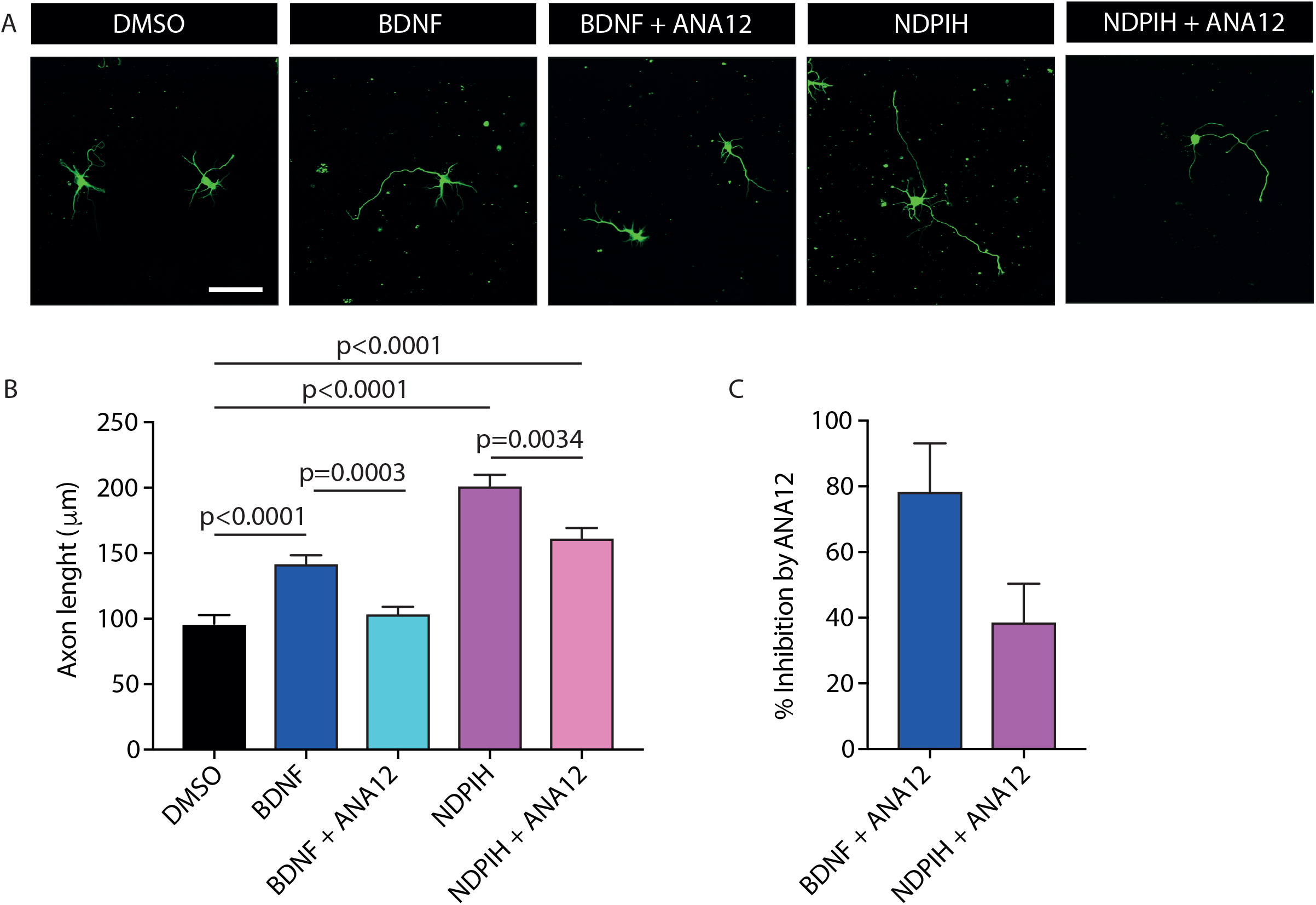
NDPIH purified from Lion Mane mushroom A2 exerts a BDNF-like neurotrophic effect in hippocampal neurons. Hippocampal neurons seeded at low density (10,000 per coverslip) to prevent paracrine effects until DIV2. Cells were then exposed to purified NDPIH (10 μg/ml) or BDNF (1nM) for 24h in the presence or absence of Trk-B inhibitor ANA12 (0.5 μM), fixed, processed for immunochemistry against β-tubulin (green) and imaged using confocal microscopy (nuclear DAPI in blue). ANA12 was added simultaneously to BDNF or NDPIH. (A) Representative images of hippocampal neurons in the indicated conditions. (B) Axon length quantification for each treatment. Axon was considered as the longest neurite. (C) Relative inhibition of axon growth done by ANA-12 on BDNF or NDPIH. Data shows mean ± SEM. n=30-60 neurons in each condition, from 3 independent neuronal preparations. One-way ANOVA, Turkey’s multiple comparison test was performed in (B). P-value is indicated when significative differences were found. Scale bar 100 µm.

Having demonstrated that *H. erinaceum* extracts have a BDNF-like activity in hippocampal cultures, we decided to test two concentrations (100 mg/kg and 250 mg/kg) of the crude extract (A1) in a rodent *in vivo* model. The potential effect of A1 on short-term memory was evaluated using the Y-maze test (Fig. 6A). Only the nootropic chemical piracetam (PC) group, a positive control known to enhance cognitive stills (68.79 ± 0.92 %, *P*^*^ < 0.05) exhibited a significant increase in alternation behavior, compared with control (vehicle) group (54.10 ± 4.78%). However, the A1-treated groups (100 mg/kg, 54.77 ± 1.41%; 250 mg/kg, 56.06 ± 2.42%) did not exhibit any spontaneous alternation behavior compared to piracetam (Fig. 6B). We used the same cohort to test novel object recognition (NORT), another form of memory. The A1-treated groups (100 mg/kg, 50.73 ± 1.97%; 250 mg/kg, 56.60 ± 4.14%) displayed significantly enhanced ability to recognize the novel object compared to the control (vehicle) group (38.59 ± 3.08%) (Fig. 6B). Furthermore, the effects of A1 groups for the novel object recognition were higher than in the piracetam group (PC, 40.88 ± 3.70%), demonstrating that A1 facilitates this type of memory (Fig. 6B). To test whether the A1 treatment elevated neurotrophins in the brain, we measured their levels by performing western blot analysis (Fig. 6C). A1 promotes a marked increase in BDNF levels at the two concentrations tested (Fig. 6C). Similarly, NGF and GDNF were also significantly increased suggesting that A1 could also promote its enhancing effects on memory via these neurotrophic factors pathways (Xiao 2016). Further, GAP-43, as a marker of axonal and synaptic growth, was also significantly increased in treated A1 groups as compared with PC group (Fig. 6C).

**Figure 6.**
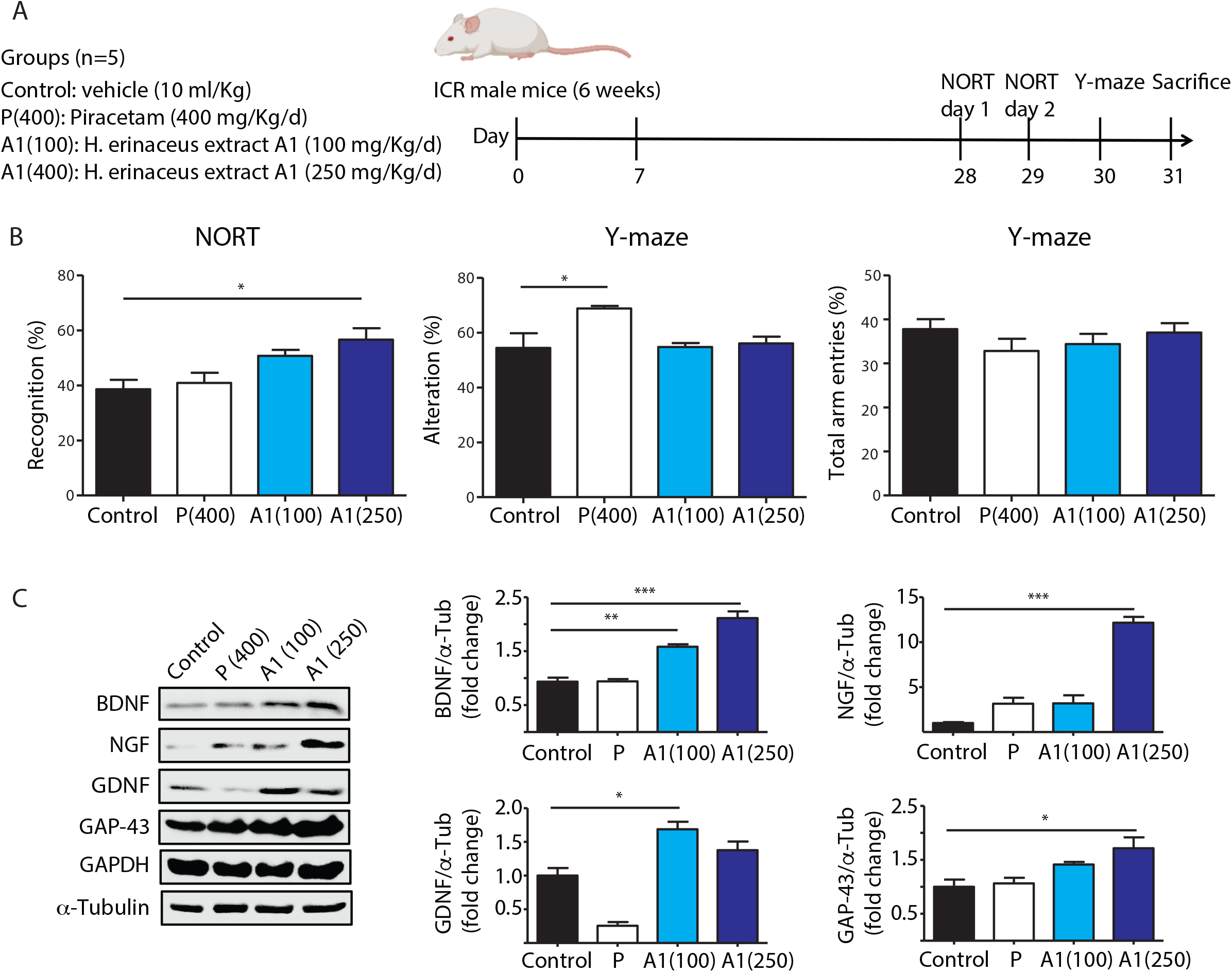
Lion Mane mushroom A1 dietary treatment improves recognition memory. (A) Mice were treated with A1 (100 or 250 mg/kg) and piracetam (PC, 400 mg/kg) as indicated in the scheme prior to the beginning of behavioral testing and treatment continued throughout the experiment. After testing, mice were sacrificed and tissue was harvested. Each sample treatment lasted a total of 3 weeks. (B) Behavioral testing. NORT test: portions of time spent exploring novel object were measured and expressed as %. Y-maze test: spontaneous alterations of number of arm entries expressed as %. All data are expressed as the mean ± SEM (n = 4 - 5). ^*^ *P* < 0.05 as compared to the normal group. (C) Representative western blots of BDNF, NGF, GDNF and GAP-43 in the whole brain. GAPDH and α-tubulin were used as loading control. All data are expressed as the mean ± S.E.M. (n = 3). ^*^*P* < 0.05, _**_*P* < 0.01 and ^***^*P* < 0.001 as compared to the control group.

Neurotrophins act by activation of ERK1/2 to induce phosphorylation of p-CREB (Mesripour *et al*. 2016). The ERK is a key signalling pathway downstream of BDNF, and alteration of the phosphorylation pattern of both CREB and ERK has been proposed as a mechanism contributing to neurodegenerative diseases (Gourley *et al*. 2008). We examined the phosphorylation of CREB and ERK among the treatment group in brain sections. Only the treated-A1 group with the highest concentration (250 mg/kg) showed significantly increased ratio of p-ERK/ERK and p-CREB/CREB, compared with the PC group (Supp Fig. 2). Overall, our findings suggest that A1 exhibit significant neurotrophic effect by modulating a range of neurotrophic pathways including NGF, BDNF, GDNF, GAP-43, ERK and CREB phosphorylation and expression in the mouse brain.

Based on our results with the *in vitro* analysis on subfractions of *H. erinaceus*, we selected hericene A to conduct the two behavioral tests because of its hydrophobic profile conducive to BBB penetration and of its significant neurotrophic activity (see Fig. 2B). Because hericene A was a purified derivative of the crude extract A1, we decided to use lower concentrations of hericene A (5 mg/Kg) to test for its ability to enhance memory function (Fig. 7A). Treatment with hericene A on the Y-maze test slightly increased (65.06 ± 0.81%) spatial memory compared to the control group (Fig. 7B). The positive control PC group exhibited a significant increase in spontaneous alternation behavior (72.91 ± 5.69 %, *P*^*^ < 0.05), compared with the control group (54.88 ± 1.48%). The role of hericene A on recognition memory was carried out using the NORT test. The hericene A groups displayed significantly enhanced ability to recognize the novel object (55.32 ± 2.54%) compared to that of normal group (41.32 ± 4.18%) (Fig. 7B). These results indicate that the administration of hericene A significantly improves recognition memory function. We also examined whether hericene A treatment increased the levels of neurotrophins. As expected, the administration of hericene A significantly increased the levels of BDNF and NGF expressions as compared with control group (Fig. 7C). This finding suggests that memory-enhancing effects of hericene A is associated with BDNF and NGF activation in the mouse brain.

**Figure 7.**
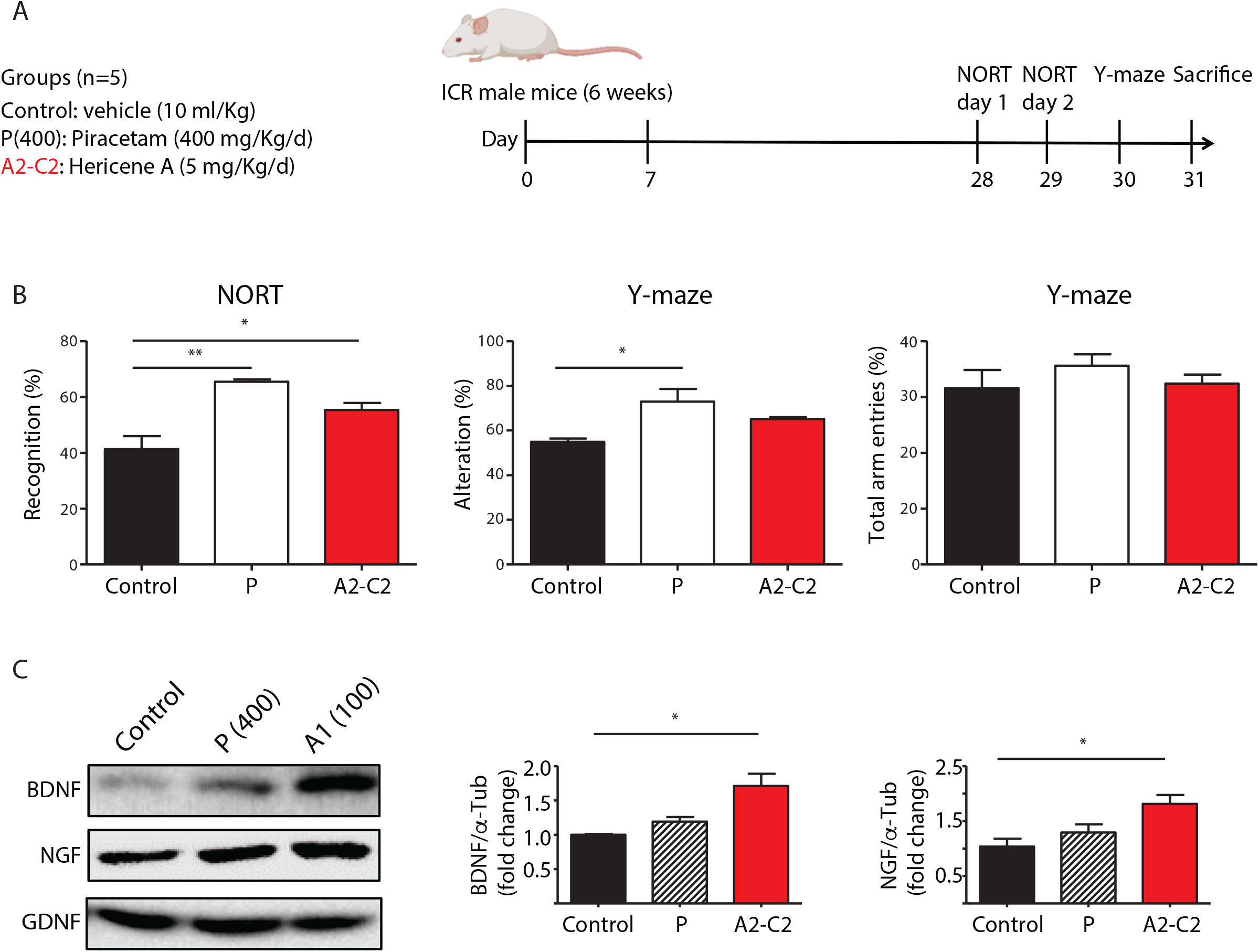
Lion Mane mushroom A2-derived hericene A dietary treatment improves recognition memory. (A) Mice were treated with hericene A (5 mg/kg) and piracetam (PC, 400 mg/kg) as indicated in the scheme prior to the beginning of behavioral testing and treatment continued throughout the experiment. After testing, mice were sacrificed and tissue was harvested. Each sample treatment lasted a total of 3 weeks. (B) Behavioral testing. NORT test: portions of time spent exploring novel object were measured and expressed as %. Y-maze test: spontaneous alterations of number of arm entries expressed as %. All data are expressed as the mean ± SEM (n = 4 - 5). ^*^ *P* < 0.05 as compared to the normal group. (C) Representative western blots of BDNF, NGF, GDNF in the whole brain. GAPDH and α-tubulin were used as loading control. All data are expressed as the mean ± S.E.M. (n = 3). ^*^*P* < 0.05 and ^**^*P* < 0.01 as compared to the control group.

## Discussion

Since the discovery of the first neurotrophin, NGF, more than 70 years ago, countless studies have demonstrated their ability to promote neurite regeneration, prevent or reverse neuronal degeneration and enhance synaptic plasticity. These, have attracted the attention of the scientific community to implement strategies through their use as treatment for a number of neurological disorders (Chao *et al*. 2006). Unfortunately, their actual therapeutic applications have been limited and the potential use of their beneficial effects remain to be exploited. Neurotrophins, for example, have poor oral bioavailability, and very low stability in serum, with half-lives in the order of minutes (Zhang *et al*. 2014). Once administered, neurotrophins have minimal BBB permeability and restricted diffusion within brain parenchyma (Pardridge 2002). In addition, their ability to activate multitude receptor signalling networks conferred pleiotropism that often resulted in undesired off-target effects such as pain, spasticity and even neurodegeneration (Constandil *et al*. 2011; Endo *et al*. 2009; Fouad *et al*. 2013; Fahnestock *et al*. 2001; Mufson *et al*. 2012; Dyck *et al*. 1997; Bergmann *et al*. 1998). As a consequence, alternative strategies to increase neurotrophin levels, improve their pharmacokinetic limitations or target specific receptors have been developed (Longo & Massa 2013). Identification of bioactive compounds derived from natural products with neurotrophic activities provided new hope in the development of sustainable therapeutical interventions.

In our study, we demonstrate that crude and purified extracts obtained from the nootropic mushroom *H. erinaceus* have BDNF-like neurotrophic activity both in cultured hippocampal neurons and in paradigm models of learning *in vivo*, leading to marked neurite outgrowth and improved memory, respectively. *H. erinaceum* mushroom characteristically contains aromatic and diterpene skeletons as active components, providing a large source of structurally related terpenoids, such as erinacines, hericenones, hericerins, hericenes, hericenols, and erinacerins. In fact, *H. erinaceus* contains 20 out of all 24 known diterpenoids (Friedman 2015). In our study, we purified four aromatic compounds from the CH_2_Cl_2_ fraction (C1 to C4) with prenyl side chains: NDPIH, hericene A, corallocin A, and 4-[3′,7′-dimethyl-2′,6′-octadienyl]-2-formyl-3-hydroxy-5-methyoxybenzylalcohol. Some of these compounds have already been isolated from *H. erinaceus*, reporting anti-diabetic and anti-cancer biological activities (Li *et al*. 2015). Importantly, we found that these four compounds have the ability to elicit a strong neurotrophic response in neuronal cultures in the absence of serum supplementation. This indicates that they have the ability to replace the initial requirement for serum trophic factors in neuronal cultures. Consistent with our present study, corallocin A (C3) was shown to induce NGF and BDNF expression in astrocytes (Wittstein *et al*. 2016). Several compounds from *H. erinaceum* have been suggested to improve brain function via NGF-related activity. Hericenones aromatic compounds, for example, stimulate NGF secretion in cultures of PC12 cells (Phan *et al*. 2014) and erinacines diterpene compounds stimulate NGF synthesis in *in vitro* and *in vivo* systems (Li *et al*. 2020; Tsai-Teng *et al*. 2016).

Neurotrophins also regulate the plasticity and growth of axons within the adult central and peripheral nervous system towards their appropriate targets (Guthrie 2007). However, the expression of many neurotrophic factors is greatly reduced within the adult central nervous system (CNS). Exogenous application of these growth-permissive neurotrophic factors has therapeutic potential through creation of favorable environments after nerve injury (Keefe *et al*. 2017). Supplementation of NGF and BDNF have been used to promote survival and axonal regrowth of injured neurons after spinal cord injury (Harvey *et al*. 2015). In this context, *in vitro* studies using total extract of *H. erinaceus* reported regenerative capabilities on peripheral neurons after laser microdissection (Ustun & Ayhan 2019). This open the door to additional studies using specific and more potent derivatives from *H. erinaceus* on their potential therapeutical action during axon regeneration.

To the best of our knowledge, our study is the first to identify a pro-BDNF activity for *H. erinaceus* and to identify hericene A as an active component for this neurotrophic function *in vitro* and *in vivo*. Our study reveals that hericene A enhances recognition memory at concentrations as low as 5 mg/kg. The medicinal mushroom *H. erinaceus* has long been known for its neurotrophic activity on the peripheral nervous system through by promoting NGF synthesis (Lai *et al*. 2013; Raman *et al*. 2015; Ustun & Ayhan 2019; Haure-Mirande *et al*. 2017). Several compounds extracted from *H. erinaceus* have been shown to protect from ageing-dependent cognitive decline in wildtype (Brandalise *et al*. 2017; Rossi *et al*. 2018) and in Alzheimer’s disease mouse models (Tsai-Teng *et al*. 2016; Mori *et al*. 2011) suggesting a yet to be defined activity on the CNS. A recent study has determined that erinacine A and hericenones C and D were capable of slowing cognitive declines in a model of frailty (Ratto *et al*. 2019). The neuroprotective effects were attributed to the ability of these molecules to elicit NGF (Mori *et al*. 2009) and BDNF synthesis (Chiu *et al*. 2018). In agreement with previous studies, we identified that polysaccharides isolated from *H. erinaceus* extracts also exert a neurotropic activity. The effects of *H. erinaceus* on recognition memory were previously shown and hypothesized to stem from promoting neurogenesis (Rossi *et al*. 2018; Brandalise *et al*. 2017).

Our study demonstrates that hericene A exhibits neuritogenesis activity and that this effect can be achieved at very low concentration both *in vitro* and *in vivo*. The highest contents of Hericene A were detected in *H. erinaceus* A1 extract, compared to the other species. To increase the production of these compounds, it might be possible to engineer new methods to improve selection of specific *H. erinaceus* strains based on their ability to produce larger quantities of hericene A derivatives. We should also provide an additional step towards hericene A-enriched *H. erinaceus*, marker compound-based standardization for sustainable use of dietary supplements and medicinal products from *H. erinaceus*. Based on our results, we suggest that the administration of hericene A from *H. erinaceu*s may be useful to ameliorate brain function and disorder-related pathologies in *in vivo* neurodegenerative disease models. Although further studies are required to test this hypothesis, our report represents the first step on exploring the potential nootrophic benefits from hericene A compounds isolated from the mushroom *H. erinaceus*.

## Acknowledgments

This work was supported by an NHMRC Senior Research Fellowship (GNT1155794), Australian Research Council LIEF Grant LE130100078 and CNG-Bio grant Funding to F.A.M. MKL was funded by Medical Research Centre program (2017R1A5A2015541) and the National Research Foundation of Korea.

