## Supplementary figures and images for "Hericerin derivatives from *Hericium erinaceus* exert BDNF-like neurotrophic activity in central hippocampal neurons and enhance memory"

### Supp Fig 1

SUPP FIGURE 1

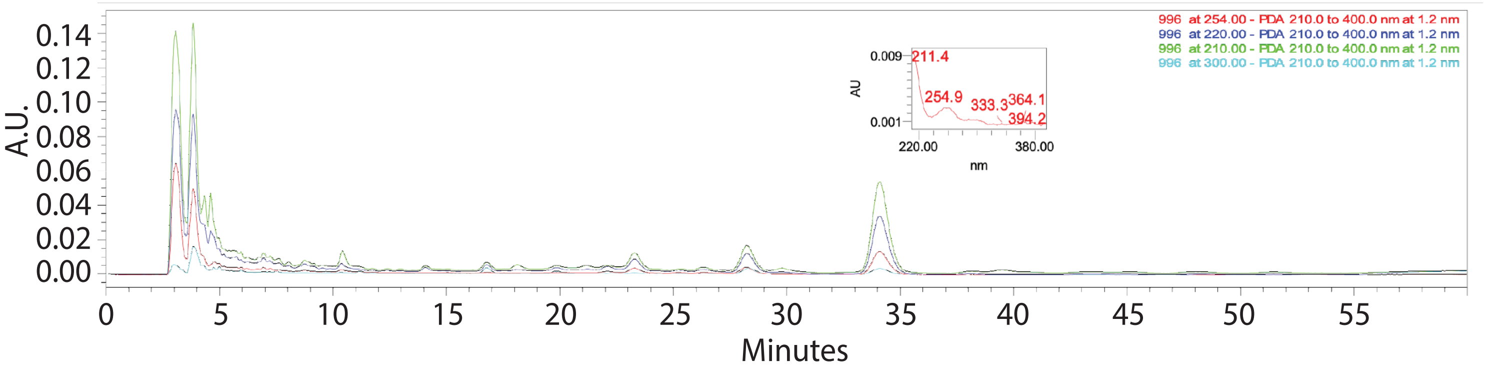

### Supp Fig 2

SUPP FIGURE 2

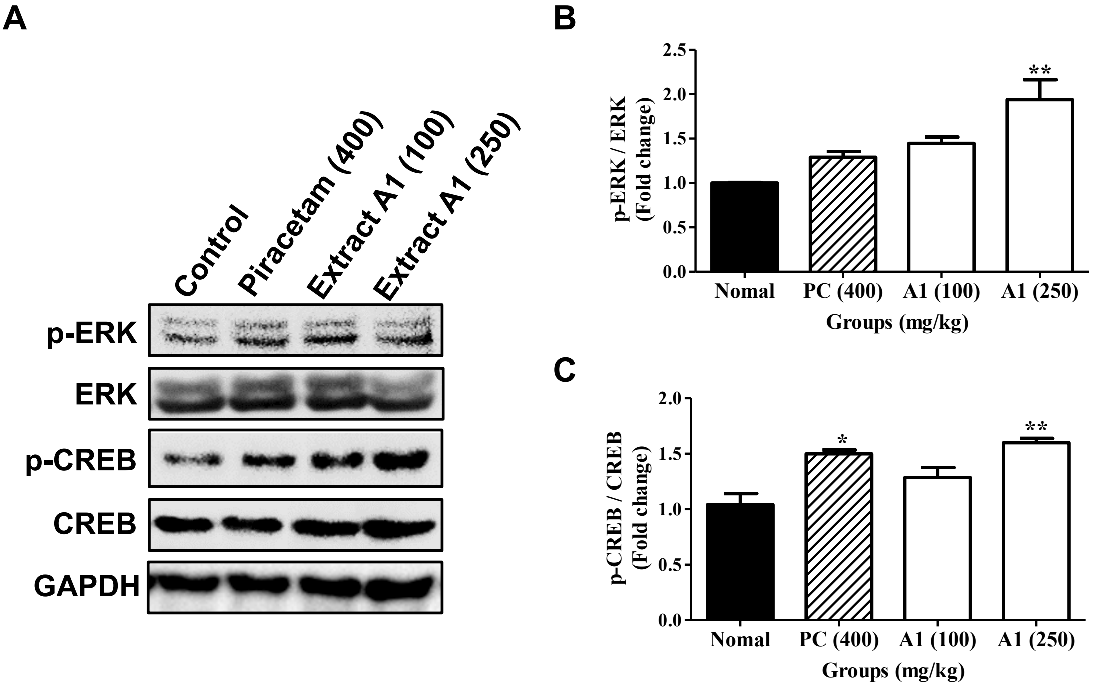
